# A resampling-based approach to share reference panels

**DOI:** 10.1101/2023.04.07.535812

**Authors:** Théo Cavinato, Simone Rubinacci, Anna-Sapfo Malaspinas, Olivier Delaneau

## Abstract

For many genome-wide association studies, imputing genotypes from a haplotype reference panel is a necessary step. Over the past 15 years, reference panels have become larger and more diverse, leading to improvements in imputation accuracy. However, the latest generation of reference panels is subject to restrictions on data sharing due to concerns about privacy, limiting their usefulness for genotype imputation. In this context, we propose RESHAPE, a method that employs a recombination Poisson process on a reference panel to simulate the genomes of hypothetical descendants after multiple generations. This data transformation helps to protect against re-identification threats and preserves important data attributes, such as linkage disequilibrium (LD) patterns and, to some degree, Identity-By-Descent (IBD) sharing, allowing for genotype imputation. Our experiments on gold standard datasets show that simulated descendants up to eight generations can serve as reference panels without significantly reducing genotype imputation accuracy. We suggest that this specific type of data anonymization could be used to generate synthetic reference panels available under less restrictive data sharing policies.

## I Introduction

Genotype imputation is the statistical estimation of missing genotypes in SNP array data, using a reference panel of sequenced individuals. Genotype imputation is ubiquitous in the field of statistical genetics and genome-wide association studies (GWAS), as it drastically increases the number of genetic variants available which helps boost association signals, identify causal variants and meta-analyze multiple cohorts ^1^. Genotype imputation predicts missing data in a target sample by considering target haplotypes as mosaics of reference haplotypes ^2–4^. The most commonly used imputation model is based on the Li and Stephens hidden Markov model ^5^ that probabilistically builds haplotype mosaics in agreement with the variable recombination rate in hotspots and coldspots observed in the human genome ^6^. In the last 10 years, the size of the reference panels used for genotype imputation has increased considerably, improving the accuracy of genotype imputation, in particular at rare variations. This has been possible thanks to the establishment of large scale projects such as the 1000 Genomes Project (1000GP) ^7^, the Haplotype Reference Consortium ^8^, the TOPMed program ^9^ and more recently the UK Biobank resource ^10–12^. However, reference panels part of national-wide biobanks, with sample sizes in the order of hundreds of thousands of genomes, comprise sample-level genetic and phenotypic data linked together, which puts strict restrictions on accessibility and data sharing and therefore prevents their wide usage for genotype imputation in other studies. In this work, we present RESHAPE (REcombine and Share HAPlotypEs), a method that enables the creation of a synthetic haplotype reference panel by simulating hypothetical descendants of reference panel samples after a user-defined number of meiosis. Such synthetic reference panel allows for high-quality genotype imputation while (i) mixing the genetic data across samples, (ii) disrupting genome-phenome links and (iii) preventing usual re-identification threats faced by pseudo-anonymised reference panels ^13^. We assessed the impact of our approach on imputation accuracy using multiple gold standard reference panels, varying the number of generations used to simulate descendants. Our approach aims to facilitate reference panel sharing by proposing an alternative dataset that is still useful for imputation and can be made available under less restrictive data sharing policies.

## II Material and methods

Through meiosis, the two haplotypes from a parent recombine, thereby creating a new haplotype that will be transmitted to the offspring. This process is repeated at each generation, involving that an individual’s haplotypes can be represented as mosaics of haplotypes of its ancestors. The method we propose in this work builds on this idea and consists in two separate steps. First, it samples recombination points in the genome based on a genetic map and a user-defined number ***K*** of simulated meiosis. Second, it recombines reference haplotypes at these positions in order to generate a given number of offspring haplotypes in which the linkage disequilibrium (LD) structure and, to some extent, Identity-By-Descent (IBD) sharing are preserved (**Fig. 1A**). The resulting synthetic reference panel is of the same size as the original, maintains the necessary information for the Li and Stephens imputation model, and disrupts the association of genotypes with individuals and therefore with any phenotypic data. Sharing a synthetic reference panel can be seen as distributing the haplotypes of relatively distant relatives of the original samples included in the reference panel.

**Fig. 1.**
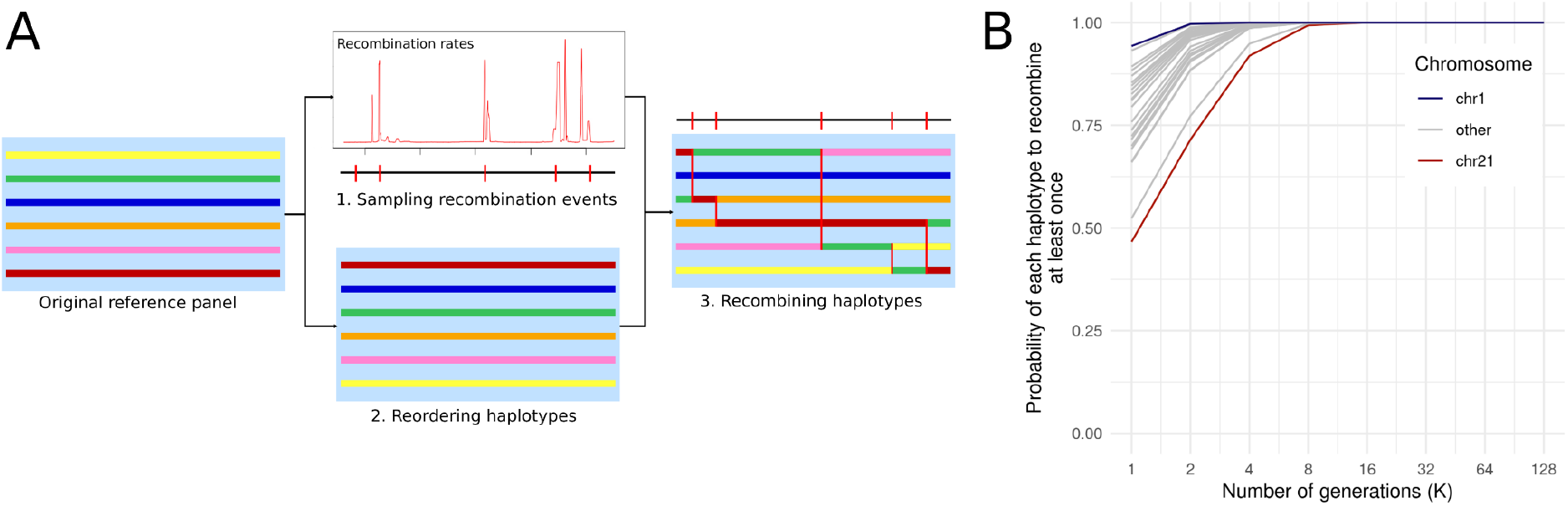
Description of the method. **A** The method takes a reference panel as input and (1) samples recombination sites using a Poisson process and a genetic map, (2) randomly reorders the haplotypes and (3) recombines the haplotypes based on the drawn recombination sites, resulting in a synthetic reference panel. **B** Probability of each haplotype of the reference panel to recombine at least once using our approach for different chromosome size in cM. Each of the 22 lines correspond to the result for a given chromosome size in cM. The blue line corresponds to the largest chromosome (chromosome 1, 286.28 cM) and the red line to the smallest chromosome (chromosome 21, 62.79 cM).

### Algorithm

Let **N** and **L** be the number of haplotypes and variant sites in the reference panel. Naive implementation of the scheme described above would involve multiple forward simulations in which a new synthetic reference panel is created at each generation. The computational cost in this case would be proportional to O(**KNL**). In our work, we propose an alternative two-steps approach which has the advantage of involving a computational cost proportional to O(**NL**), essentially independent of **K**.

In the first step, we model the total number of recombination events in a specific genomic region as a Poisson distribution with **λ** equal to the average number of recombination sites expected in that region i.e. the size of this region in Morgan times (**N/2)K**, to account for the number of generation simulated and the number of haplotypes in the dataset. Thus, we can simulate genomic positions at which recombination events occur along a chromosome using a Poisson process, which consists of drawing genetic distances in centiMorgan (cM) from the inverse cumulative distribution function (CDF^-1^) of a Poisson distribution. This results in an array **R = {r**_**1**_, **…, r**_**M**_**}** of genomic positions with **M** being the total number of recombination events drawn. Finally, all these genetic positions in cM are converted into base pairs (bp) using the genetic maps provided as input.

In the second step, we start by shuffling the indexes of the reference haplotypes to establish an initial random order. Then, we stream the haplotype data in an output file according to this new order for all haplotypes at every site until we reach the first recombination event in **R**. When this happens, we update the current ordering by permuting (recombining) two randomly sampled haplotypes (with replacement) and we carry on streaming the data until we reach the next recombination event. We repeat this procedure until the end of the chromosome, permuting two haplotypes each time we encounter another recombination event. The synthetic reference panel comprises the same number of haplotypes and variant sites as the original dataset. This approach has two important properties. First, the data transformation is straightforward to reverse. Indeed, we can easily regenerate the full sequence of events leading to the synthetic data knowing the genetic map, the value of K and the seed of the random number generator and therefore restore the data to its original state. Second, some haplotypes may not recombine. The probability that a given haplotype recombines at least once can be derived from a Poisson distribution and increases with K and the length in cM of the chromosome (**Fig. 1B**). Thus, with K>=8, even the smallest chromosome (chromosome 21, 62.79 cM) has a probability of recombining at least once that exceeds 0.99.

We implemented this procedure and its reverse in a C++ software called **RESHAPE** (**RE**combine and **S**hare **HAP**lotyp**E**s). By providing it with a genetic map and the VCF/BCF of a reference panel, **RESHAPE** outputs a VCF/BCF of the same size containing descendant haplotypes.

### Benchmark

In our experiments, we used phased data from two well-known reference panels of haplotypes: (i) the 1000GP ^7^ comprising data for 2’504 samples and (ii) the the first release of the phased UK Biobank which consists of 147’754 samples ^10–12^. In each dataset, we filtered out monomorphic and multi-allelic variants and only retained data on chromosome 20. Number of samples, sample ancestries and number of variant sites can be found in **Supplementary Table 1**. We used the genetic map of chromosome 20 derived from the HapMap project ^14^. For each genetic position of a reference panel not present in the genetic map, **RESHAPE** automatically infer the corresponding position in cM using linear interpolation. As a null, we also generated additional genetic maps by keeping the HapMap size of the chromosome in cM but assuming a constant recombination rate. We used **RESHAPE** to generate multiple synthetic reference panels using K=1, 2, 4, 8, 16, 32 and 128. We also used the original reference panels that we called K=0.

To quantify the depletion of LD due to the introduction of recombination events in the data, we used LD scores. The LD score of a given variant is defined here as the sum of r^2^ between this variant and all other variants in its vicinity (+/- 100Kb). Global changes in LD scores were primarily assessed using Pearson’s correlation coefficient between the LD-scores of the original and the synthetic reference panels. We complemented this by fitting linear regression models using the LD-scores of the synthetic and original panels as explanatory and dependent variables, respectively. With a r² close to 1, any deviation from 1 in the regression slope (β) would reflect a global increase (β>1) or decrease (β<1) in LD scores.

To assess the reduction in imputation accuracy, we mimicked SNP array data by using genomes sequenced at high-coverage. We masked genotypes at markers not present in three different SNP arrays, then we imputed back these missing genotypes using the original and synthetic reference panels with BEAGLE v.5.3 ^2^ and finally compared the imputed genotypes to the high-coverage ones. We performed two imputation experiments based on the high-coverage sequencing data for 2,504 samples from the 1000GP ^15^ and for 147’754 samples from the UK Biobank ^10–12^. In each dataset, we extracted a small number of samples to act as target samples and used the remaining ones as reference panel data, synthetic or not (**Supplementary Table 1**). We simulated two types of SNP arrays for experiments on the 1000GP: the Illumina HumanOmni 2.5 array and Illumina Global Screening Array. For the experiments on the UK Biobank, we only used the UKB Axiom array. The concordance between true and imputed genotype calls was computed using GLIMPSE_concordance v.1.1.1 ^4^.

## Results

We investigated the effect of the number of generations (*K*) onto LD-scores in two populations: the European and African samples of the 1000GP. Overall, we find that our recombination approach has low impact on LD-scores, even with high number of generations (i.e. *K*=128), and maintains high correlation levels with the original LD-scores (**Fig. 2A**). We however observed a general decline in LD-scores, as expected by the introduction of new recombination events in the data (**Fig. 2B**). This decline is far more pronounced when using a constant-rate genetic map than with a HapMap genetic map, consistent with the more frequent occurrence of highly LD disruptive recombination events in recombination coldspots. Of note, LD scores for the African population are slightly less affected than for the European population, likely due to the overall lower LD in this population. Finally, closer inspection of pairwise LD values revealed that fine LD structure is remarkably well preserved even when using a high number of generations (**Fig. 2C**).

**Fig. 2.**
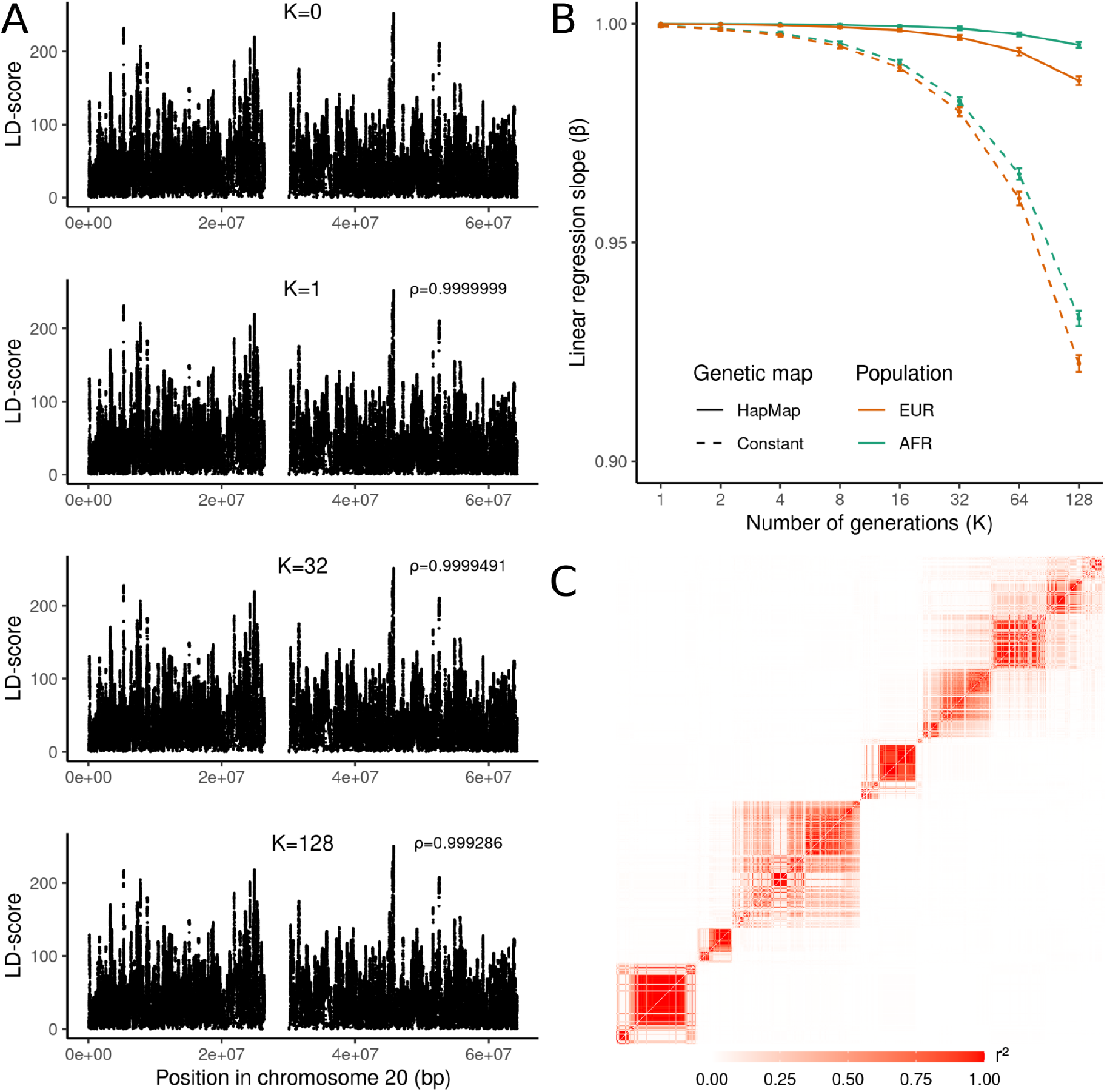
LD-score of synthetic haplotypes. **A** Average LD-score through 50 replicates of the European reference panel generated using *K*=0, 1, 32 and 128. The value **ρ** is the Pearsons’s correlation coefficient between LD-scores obtained for *K*=0 and LD-scores obtained for *K*=1, *K*=32 and *K*=128. **B** Average linear-regression slope (β) between the original reference panel and synthetic reference panels through 50 replicates (y-axis) for different values of *K* (x-axis). Colors correspond to the population analyzed (orange for the Europeans (EUR) and green for the Africans (AFR)). The error bars correspond to the mean of the 50 replicates +/- the standard deviation. **C** Correlation (r²) between SNPs in a 300 Kbp long region of chromosome 20. Values above the diagonal were obtained from the original European reference panel and values under the diagonal were obtained from a synthetic European reference panel generated using *K*=128.

We also assessed the extent to which recombining haplotypes in a reference panel affects genotype imputation. To do so, we imputed genotypes in 52 individuals from the 1000GP (2 individuals from each population) using the remaining 2’452 samples as reference panels, either recombined at different levels or not. Overall, we find that our recombination approach with up to *K*=8 has low impact on imputation and maintains a decrease in imputation accuracy below 0.01, regardless of the SNP array used (**Fig3 A-B**). Further recombining the data up to *K*=128 leads to a moderate decrease of 0.07 in imputation accuracy. Importantly, we find that rare variants seem to be more affected by the recombination procedure than common variants, for which almost no loss of accuracy is observed (**Fig3 C-D**). To further investigate this effect on rare variants, we used the sequencing data for 147’754 UK Biobank samples in our imputation experiments. We split the dataset in 1’000 target and 146’754 reference samples, generated synthetic reference panels with various numbers of generations and used the resulting data to impute the target samples. We find similar patterns than for the 1000GP data: common variants are well imputed regardless of the number of generations conversely to rare variants that are more affected **(Fig. 4)**. The largest drops in imputation accuracy are reached for extremely rare variants with a MAF < 0.005% (1 haplotype out of 20,000 carries a copy of the minor allele): 0.03, 0.10 and 0.32 for *K=*8, *K*=32 and *K*=128, respectively. Two important observations can be made here. First, the drop in imputation accuracy remains below 0.05 for a number of generations up to *K*=8, suggesting that *K*=8 maintains high imputation accuracy even in very large reference panels. Second, the important loss of accuracy involved by *K*=128 still provides better imputation accuracy than the publicly available and not-recombined HRC panel ^8^.

**Fig. 3.**
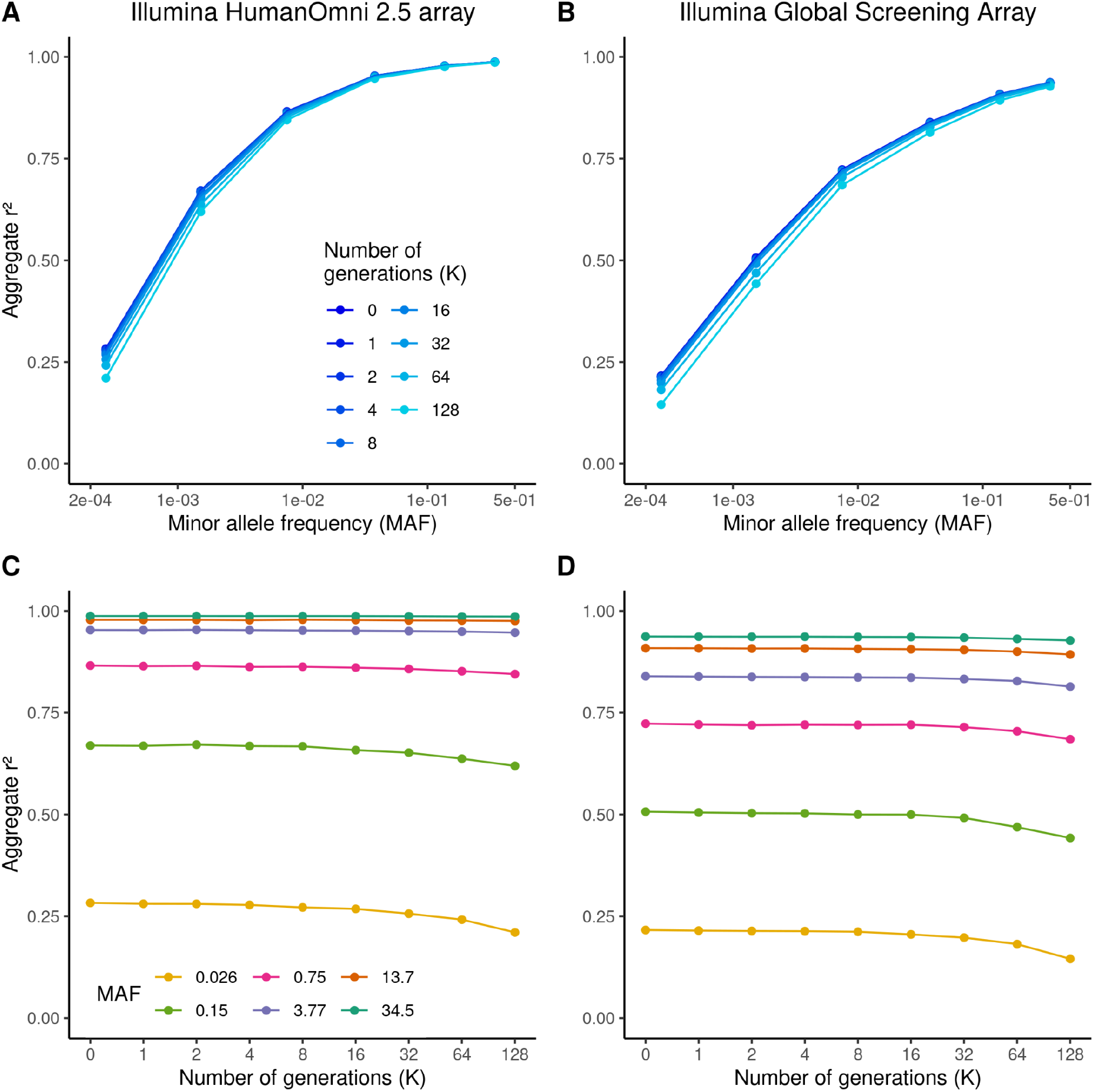
Imputation accuracy on synthetic haplotypes from the 1000GP. Aggregate r² depending on the number of generations used to recombine the haplotypes. **A-B** Shades of blue corresponds to the values of *K* used to recombine the haplotypes. **C-D** Colors correspond to MAF bins. Results with the Illumina HumanOmni 2.5 array are displayed on the left side of the panel and results with the Illumina Global Screening Array on the right side.

**Fig. 4.**
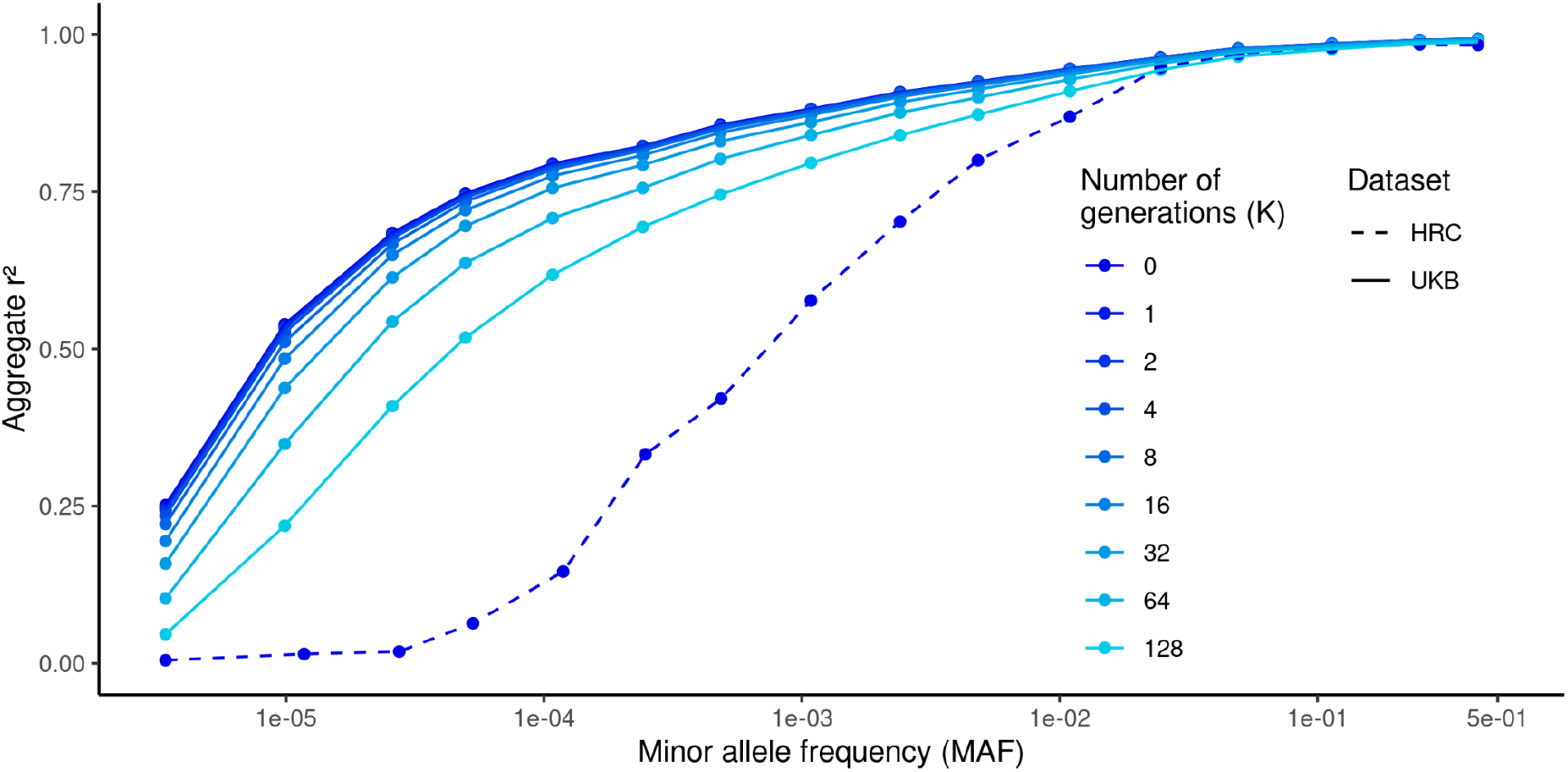
Imputation accuracy on synthetic haplotypes from the UK Biobank. Aggregate r² depending on the number of generations used to recombine the haplotypes (shades of blue). Dashed and solid lines distinguish results obtained with the HRC from the one obtained with the UK Biobank.

## Discussion

We presented an approach to remove genome-phenome links in a reference panel while maintaining key features necessary for genotype imputation. Briefly, our approach consists of sampling haplotype fragments from a reference panel to create synthetic haplotypes, which can be seen as a mosaic of the original haplotypes. The synthetic haplotypes have a high chance to be fragmented at positions with a high recombination rate, which explains the low impact of our approach on LD-structure. The IBD segments originally shared between the reference panel and a target sample are therefore partially conserved, and can thus be used by imputation methods for inferring missing genotypes. Similar approaches consisting of sampling real genetic data to generate *in silico* datasets have been implemented in HAPGEN ^16,17^ and HAPNEST ^18^: tools which simulate individual-level genotypic and phenotypic data to assess the effect of different genotyping chips on GWAS power or to compare Polygenic Risk Scoring (PRS) methods.

There is a trade-off between the level at which we recombine the reference haplotypes and the conservation of LD-structure and imputation accuracy: these two properties decay with *K*, but the higher the *K* and the higher the chance of all haplotypes to recombine at least once. We have shown that *K*=8 is a reasonable solution since (i) the probability of all haplotypes to recombine at least once is above 0.99, (ii) the LD-structure is highly conserved and (iii) imputation accuracy is comparable to imputation accuracy obtained with the original reference panel. These conclusions hold for synthetic reference panels generated using a HapMap genetic map. Constant-rate genetic maps, in addition to strongly decreasing LD-structures, could facilitate the identification of the sampled recombination sites and therefore may enable the reconstruction of the original haplotypes.

A synthetic reference panel generated with a sufficiently high number of generations and a real genetic map may be easier to share to the scientific community for imputation purposes. Two models can be considered here: either making the synthetic reference panel accessible in a centralized imputation server or directly sharing it with users for local imputation. These two solutions increase the genetic privacy of the reference panel’s data providers and of the target sample to be imputed, respectively. The absence of genome-phenome link in the synthetic reference panel prevents certain analyses such as genome-wide association studies or polygenic risk scores from being performed on the data. However, other analyses can still be applied to the synthetic reference panel, as long as they require data properties that are conserved, such as allele frequencies and LD structure.

This data transformation is reversible under certain conditions. First, the original reference panel can be easily restored from the synthetic panel using the same parameters that were employed to build it. In case these parameters are not available, recovering the original data may be possible if additional pieces of information are available for the original samples. One approach could involve a leak of phenotypic data. In this case, this would entail finding the mosaics of synthetic haplotypes that best explain (based on PRS) the known phenotypes. However, we anticipate that this problem would be extremely difficult to solve in practice without any guarantee of restoring the original data. Another possibility relies on the availability of SNP array data for the original samples. In this scenario, imputing the SNP array data using the synthetic reference panel should produce genetic data close to the original panel. However, imputing the SNP array data with another large-scale reference panel would lead to a similar outcome.

Removing genome-phenome links goes one step further than anonymization against re-identification threats in reference panels. In a reference panel where no information is linked to each pair of haplotypes, one could still infer phenotypes from an haplotype pair and deduce the identity of its donor ^13,19^. Recombining haplotype fragments disrupt the phenome carried by the original haplotypes, thereby avoiding such re-identification approaches. Nonetheless, we anticipate that it would be possible to use haplotype sharing techniques to test for the existence of a particular individual or a relative in a synthetic reference panel. Such an approach would require holding genotype data from an individual present in the reference panel.

With the availability of modern reference panels, several new challenges need to be addressed using different strategies and approaches. One such approach is meta-imputation ^20^, which aims to leverage multiple imputation servers based on different reference panels to accurately impute target samples. Another approach is imputation using homomorphic encryption ^21^, which aims to protect sensitive data on the user side by encrypting target genotypes before sending them to imputation servers. However, this approach is still restricted to Linear Models which are known to perform poorly on rare variants; the main benefit of using large reference panels. To complement this toolbox, we suggest a technique for generating a synthetic reference panel that retains imputation information while removing the original genome-phenome association. This could be advantageous for both centralized and local imputation servers, as the synthetic panels would no longer contain the most sensitive information, potentially increasing the willingness of large consortia to share their reference data for imputation purposes.

## Acknowledgments

This work was funded by a Swiss National Science Foundation project grant 373 (PP00P3_176977) conducted under UK Biobank project 66995. The New York Genome Center 1000 Genomes data were generated at the New York Genome Center with funds provided by a National Human Genome Research Institute grant no. 3UM1HG008901–03S1. We thank Jared O’Connell and Suyash Shringarpure from 23andMe for their insightful comments.

## Author Contributions

T.C., S.R. and O.D. designed the study. T.C. developed the algorithms and wrote the software. T.C. performed the LD experiments and the imputation experiments using the 1000GP. S.R. performed the imputation experiments using the UK Biobank. O.D. and A.S.M. supervised the project. T.C. and O.D. wrote the paper. All authors reviewed the final manuscript.

## Data and code availability

RESHAPE source code is available with MIT license from https://github.com/TheoCavinato/RESHAPE. The 1000 Genomes Project phase 3 dataset sequenced at high coverage by the New York Genome Center is available on the European Nucleotide Archive (ENA) under accession no. PRJEB31736, the International Genome Sample Resource (IGSR) data portal and the University of Michigan school of public health ftp site (URL: ftp://share.sph.umich.edu/1000g-high-coverage/freeze9/phased/). The publicly available subset of the HRC dataset is available from the European Genome-phenome Archive at the European Bioinformatics Institute (EBI) under accession no. EGAS00001001710. The UK Biobank genetic data are available under restricted access. Access can be obtained by application via the UK Biobank Access Management System (URL: https://www.ukbiobank.ac.uk/).

