## Supplementary material for "A resampling-based approach to share reference panels"

<table>
<tr><th>Dataset</th><th>#Samples</th><th>#Variants after filtering</th><th>#Samples in reference panel</th><th>#Target samples</th><th>SNP array</th></tr>
<tr><td>European (EUR) samples from the 1000GP</td><td>503</td><td>110’708</td><td colspan="3" rowspan="2">These datasets were not use in imputation analysis</td></tr>
<tr><td>African (AFR) samples from the 1000GP</td><td>661</td><td>138’699</td></tr>
<tr><td rowspan="2">1000GP</td><td rowspan="2">2’504</td><td rowspan="2">2’116’846</td><td rowspan="2">2’452</td><td rowspan="2">52</td><td>Illumina Global Screening Array</td></tr>
<tr><td>Illumina Human Omni 2.5 array</td></tr>
<tr><td>UK Biobank</td><td>147,754</td><td>13’677’164</td><td>146’754</td><td>1’000</td><td>UKB Axiom Array</td></tr>
</table>

**Supplementary table 1.** Datasets, number of samples, variants after filtering, samples in the reference panel, target samples and names of the SNP-arrays used for each analysis
